# MapGL: Inferring evolutionary gain and loss of short genomic sequence features by phylogenetic maximum parsimony

**DOI:** 10.1101/827907

**Authors:** Adam G Diehl, Alan P Boyle

## Abstract

Comparative genomics studies are growing in number partly because of their unique ability to provide insight into shared and divergent biology between species. Of particular interest is the use of phylogenetic methods to infer the evolutionary history of cis-regulatory sequence features, which contribute strongly to phenotypic divergence and are frequently gained and lost in eutherian genomes. Understanding the mechanisms by which cis-regulatory element turnover generate emergent phenotypes is crucial to our understanding of adaptive evolution. Ancestral reconstruction methods can place species-specific cis-regulatory features in their evolutionary context, thus increasing our understanding of the process of regulatory sequence turnover. However, applying these methods to gain and loss of cis-regulatory features currently requires complex workflows which represent a potential barrier to widespread adoption by a broad scientific community. MapGL simplifies phylogenetic inference of the evolutionary history of short genomic sequence features by combining the necessary steps into a single piece of software with a simple set of inputs and outputs.

## Introduction

Comparative genomics uses sequence-level differences between species to gain insights into how genomes function and evolve (1). According to Google Scholar, published comparative genomics studies have increased every year since 2009. These studies rely on the ability to detect and assign provenance to lineage-specific sequence variations at both the nucleotide level and at the level of larger-scale sequence insertions and deletions (indels). While species-specific indels are easily visible as gaps in pairwise alignments, mere gap presence tells us nothing about the evolutionary process that caused the gap – i.e., whether it resulted from a species-specific sequence gain or from loss of an ancestral sequence element. Understanding the mechanisms driving sequence divergence and their relationships to natural selection requires the ability to discern between these events. Ancestral reconstruction is a phylogenetic method by which observed states in outgroup species are used to infer the state in the most-recent common ancestor (MRCA) (2). This inferred ancestral state can then be used to predict the evolutionary events leading to an indel. These methods can place indels in their evolutionary context, allowing much greater precision in hypothesis generation regarding the causes, underlying mechanisms, and downstream effects of cis-regulatory turnover.

However, ancestral reconstruction is a complex process and, while tools exist to reconstruct ancestral protein and DNA sequences (3–5) and ancestral genomes (6), no published tool exists to infer evolutionary gain and loss of short genomic sequence features. Doing so has historically relied on complex, ad-hoc workflows involving multiple mapping steps to target and outgroup genomes and subsequent analysis with a various published genomics software and custom scripts (see (7) for example). This complexity is exacerbated by the need to use multiple outgroup species in order to ensure the reliability of inferences (8); this comes at the cost of increases in the numbers of input and intermediate files, alignment and post-processing steps, the amount of disk space required, and the overall time needed to set up and run the analysis. These limitations represent a significant barrier to use of these methods by non-specialists, thus preventing widespread application by a broad scientific community. MapGL addresses this problem by combining all mapping and phylogenetic analysis steps into a single program with simple inputs and easily interpreted outputs.

## Design and Implementation

MapGL applies a simple phylogenetic algorithm to infer the evolutionary history of short genomic sequence features from a query genome relative to a target genome (Fig. 1A). For each query feature, an initial mapping step to the target genome determines whether an orthologous sequence exists. If so, the sequence is labeled as an ortholog and written to output. Otherwise, the ancestral state is inferred by projecting data observed from the query and target species onto a phylogeny describing the evolutionary relationships between all present-day and ancestral species (Fig. 1B). We infer the ancestral state by following the maximum parsimony principle in which we seek to minimize the total number of gain/loss events required to explain the pattern of sequence presence/absence in the observed data (9) (Fig. 1C). Query sequences are mapped to each outgroup and state labels recorded at the corresponding leaf nodes: “1” if an orthologous sequence exists or “0” if not. A post-order traversal is then performed to infer states at all internal nodes, choosing the most-frequent state observed among child nodes, or storing the set of all uniquely-observed symbols in case of ties. The inferred state at the root node, representing the MRCA, is returned and used to infer whether sequences were gained on the branch leading to the query species (i.e., were absent in the MRCA) or lost on the branch leading to the target species (i.e., were present in the MRCA) (Fig. 2A-B). A copy of the phylogeny with the outermost outgroup pruned (Fig. 1D) is used to resolve cases in which the root node label is ambiguous using the full phylogeny (Fig. 2C-D).

**Fig. 1:**
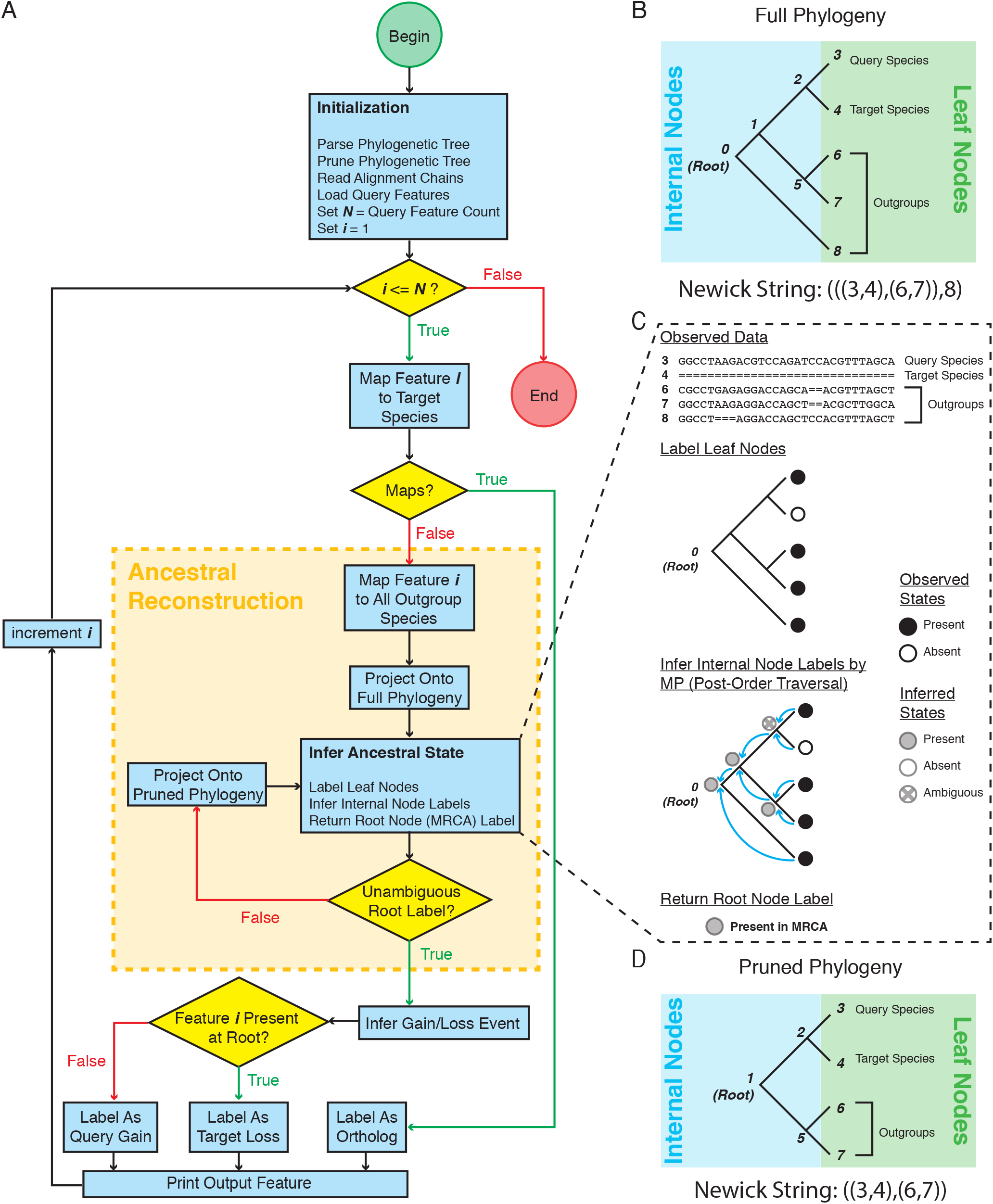
The mapGL algorithm. **(A)** Schematic outline for the mapGL algorithm. After initialization, the algorithm loops over query features, performing an initial mapping step against the target species. If the feature maps to the target species, it is labelled as an ortholog and written to output. If not, it enters the ancestral reconstruction stage. The feature is then mapped to each outgroup species in the full phylogeny and the corresponding leaves are labelled to indicate presence or absence. Internal labels are inferred based on the patterns observed at the leaf nodes (see Figure 2). If the root state cannot be inferred unambiguously, the process is repeated using the pruned phylogeny. Gain and loss events can then be inferred based on whether a feature is present at the root of the tree. The labelled feature is then written to output. This process is repeated until all query features are labelled. **(B)** Full phylogenetic tree describing evolutionary relationships between the query and target species (nodes 3 and 4) plus three outgroups (nodes 6-8). Query, target, and outgroup species occupy the leaf nodes of the tree. These are the only species for which we can directly observe sequence presence/absence. Internal nodes (0, 1, 2, and 5) represent ancestral species. **(C)** Since we cannot observe internal sequences directly, we must infer sequence presence/absence based on present-day observations from the leaf species. The core step of the ancestral reconstruction stage involves labelling all leaf nodes with their observed states and performing a post-order tree traversal to infer the states at internal nodes following the principle of maximum-parsimony (MP). The most-recent common ancestor (MRCA) occupies the root node (node 0), and the inferred state at this node is returned and used to predict whether query-specific sequences were gained in the query genome or lost from the target genome (see figure 2 (A-B) for example). **(D)** In the initialization stage of the algorithm, a copy of the full phylogeny is pruned to remove the outermost outgroup (node 8) and rerooted at the first internal node (node 1) up from the original root (node 0). This pruned phylogeny is used to resolve cases in which the root state is ambiguous (see figure 2 (CD) for example).

**Fig. 2:**
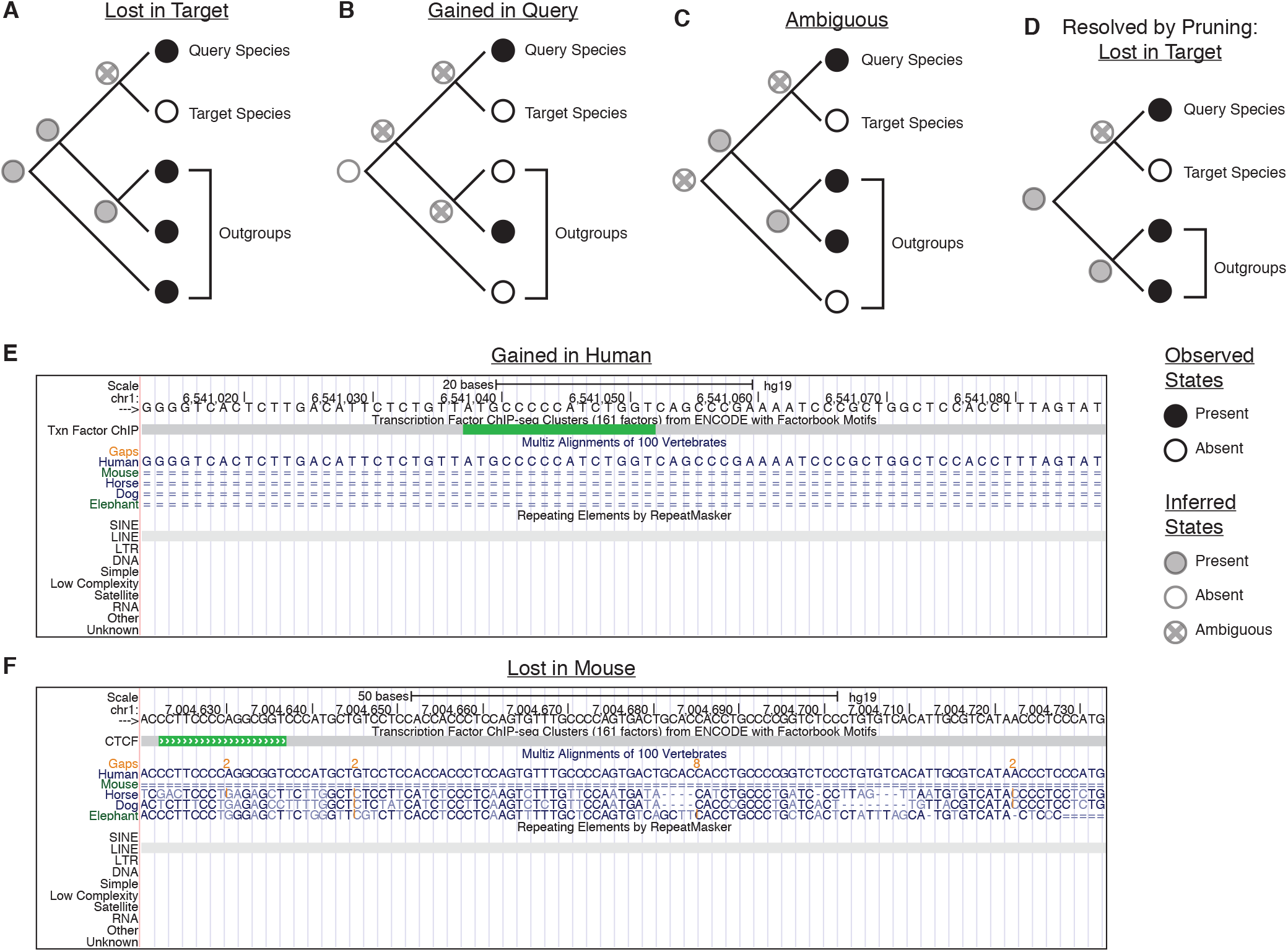
Example gain and loss inferences based on ancestral state reconstruction. **(A)** Example sequence loss in the target species based on presence in the most-recent common ancestor (MRCA), at the root of the tree. **(B)** Example sequence gain in the query species. Sequence absence is the most parsimonious inferred ancestral state as it invokes two independent sequence gains rather than three independent losses necessary to explain presence of an ancestral sequence. **(C)** The ancestral state cannot be unambiguously inferred based on the observed data. **(D)** The phylogeny in C is resolved by pruning the outermost outgroup and rerooting the tree. The inferred ancestral state is presence of an orthologous sequence and sequence loss in the target species is called. **(E)** UCSC Genome Browser track for a region in the human genome (hg19 build) labelled by MapGL as a human-specific gain. The region contains a CTCF binding annotation residing in a primate-specific insertion of an L2a LINE transposable element. **(F)** UCSC Genome Browser track for a region in the human genome labelled by MapGL as a mouse-specific loss. This region also contains a CTCF binding site residing within an L2a LINE element, but one that was inserted prior to divergence of Atlantogenata and Boreoeutheria species.

MapGL takes as inputs a set of genomic query features in BED format, a set of liftover chains (10) corresponding to the target and outgroup species, and a Newick tree describing the full phylogeny. The program is self-documenting and ships with a small example dataset. MapGL can be tuned through several command-line options. The most important of these are –threshold (−t) and –gap (−g). The −t <FLOAT> option specifies the amount of overlap required between query features and target chains to call a match as a function of the query element length. The default value of 0.0 is equivalent to requiring a single-base overlap while a value of 1 would enforce only full-length alignments. The −g <INT> option specifies the tolerance for gaps and takes an integer specifying the maximum tolerable gap length, in nucleotides, as its argument. The default value of −1 indicates that gaps of any length are allowed. The default values maximize the algorithm’s ability to find orthologous sequences, which leads to conservative gain and loss predictions. However, these values may lead to erroneous conservation predictions in cases where aligned features contain embedded indels and may be overly conservative when comparing closely-related genomes. Thus, these options may require tuning if the evolutionary distance between query and target species is very short or if the objective is to rigorously identify conserved features. Nonetheless, as the prototypical use of this program is to identify whole sequence features that have undergone species-specific gain or loss, the default values are likely reasonable for most users.

MapGL is based on bnMapper (11), which maps genomic sequences across species based on liftover chains (10). We retained the core mapping steps from bnMapper with one modification: while bnMapper drops alignments that are split across chains, we keep the longest alignment among all chains represented. We extended this framework by adding the ability to map against multiple liftover chains to incorporate target and outgroup species, and added the ancestral reconstruction steps outlined above. Tree parsing and phylogenetic methods were adapted from the python-newick package (https://pypi.org/project/newick/). MapGL is compatible with MacOS and most Unix-like operating systems, and is available for installation through PyPI and Bioconda, with source code available from the GitHub repository (https://github.com/adadiehl/mapGL).

## Results

We have applied MapGL to classify species-specific transcription factor binding sites and chromatin loop anchors in human and mouse as evolutionary gains or losses (12). Two representative human-gain and mouse-loss events clearly show that significant differences in evolutionary history can exist even when similar functional annotations are present (Fig. 2E-F). Both regions contain human-specific CTCF binding sites that are derived from L2a retrotransposon insertions. However, these two features have very different evolutionary histories: the feature shown in Fig. 2E originated from an L2a insertion that occurred after primate-rodent divergence while the region in Fig. 2F originated from an L2a insertion that occurred very early in placental mammal evolution, prior to the divergence of Atlantogenata and Boreoeutheria. Based solely on the presence of a gap in the mouse sequence, both these features may have been identified as either a human-specific gain or a mouse-specific loss, representing very different evolutionary scenarios with distinct interpretations in regards to regulatory innovation. By incorporating additional outgroup data and reconstructing the most-parsimonious ancestral state, MapGL allowed these possibilities to be resolved, placing each element in its proper evolutionary context.

## Discussion

Comparative genomics relies on the ability to infer the creative mechanisms and downstream effects of lineage-specific sequence divergence. However, even sequences with similar properties can have very different evolutionary histories. This ambiguity compromises our ability to determine how lineage-specific gain and loss of short sequence features relates to conserved and divergent functions. Placing species-specific sequence features in their proper evolutionary context enables a deeper understanding of functional divergence, but requires application of phylogenetic methods that have not been previously integrated into a single piece of software. MapGL offers a simple method to infer the evolutionary history of short sequence features, opening these methods to a much broader user base.

## Availability

MapGL is available as a Python package through PyPi, and Bioconda, with source code available at https://github.com/adadiehl/mapGL.

## Acknowledgements

We would like to thank members of the Boyle lab for critical reading and suggestions on the manuscript.

